# Endurance Exercise to Improve Physical Function in Adult and Older Mice: High Intensity Interval Training (HIIT) versus Voluntary Wheel Running (VWR)

**DOI:** 10.1101/2021.05.20.445050

**Authors:** Megan L. Pajski, Chris Byrd, Nainika Nandigama, Emily Seguin, Anna Seguin, Alyssa Fennell, Ted G. Graber

## Abstract

With age comes a gradual decline in physical function and exercise capacity, concurrent with a progressive propensity for development of sarcopenia /(age-related loss of muscle mass and strength) and frailty (inability of body to thrive and maintain homeostasis). Prior research has demonstrated that exercise, while not a cure, can help mitigate sarcopenia/frailty and restore functional capacity. Reliable, validated, pre-clinical models are necessary to elucidate the underlying molecular mechanisms at the intersection of age, exercise, and functional decline. In this study, we hypothesized that endurance exercise programs mimicking typical human exercise protocols would improve physical function in both adult and older adult mice. Furthermore, our secondary hypothesis was that older mice would receive less benefit from a similar volume of exercise than adult mice. To test these hypotheses, we randomly assigned (with some selection criteria) male C57BL/6 mice either at adult ages during the study (n=24, designated 10m, aged 6 months to 10 months) or at older adult ages (n=18, designated 26m, aged 22 months to 26 months) to a 13-week program of voluntary wheel running (VWR, group termed RUN) or high intensity interval training (HIIT), with an additional 10m sedentary control (CON). The functional aptitude of each mouse was determined pre- and post-training using our composite CFAB (comprehensive functional assessment battery) scoring system consisting of voluntary wheel running (volitional exercise and activity rate), treadmill (endurance), rotarod (overall motor function), grip meter (forelimb strength), and inverted cling (whole body strength/endurance). To measure sarcopenia, we tracked body mass and body composition changes with EchoMRI, and measured muscle wet mass post-training.

## Introduction

Sarcopenia, the age-related loss of muscle mass and strength, has long been recognized as a significant factor in quality of life among aged individuals (Tzankoff 1977, Rosenberg 1997). Studies show that after the age of fifty, muscle mass decreases by 1–2% annually (Quittan 2016) while muscular strength can decrease 12– 15% every ten years (Larsson 1983). One of the major consequences of this loss is reduced functional ability (Visser 2005, Rolland, et al. 2008, Schaap 2013), which contributes to overall frailty and can lead to a gradual loss of the ability to perform activities of daily living (ADLs), leading to loss of independence, onset of disability, and increased mortality (Marzetti et al. 2017; Reeves et al. 2004; Parks et al. 2012; Liu et al. 2014). While some of this age-related muscle loss is due to reduced physical activity and dietary protein absorption, other possible contributing factors are changes in the dynamics and regulation of inflammatory, signaling, and transcriptional processes (Graber 2015; Strasser 2007; Rolland 2008).

As early as 1991, it has been recognized that individuals and their organ systems experience differential aging, clearly indicating that environmental factors influence the extent of biological age relative to chronological age (Collier 1991). There has been particular interest in understanding the role that exercise can play in slowing the progression of sarcopenia. Low muscle mass correlates to weakness in older adults (Newman 2003), while weakness has a strong positive correlation with decreased function (Schaap 2013) and mobility (Visser 2005). Additionally, inactivity of any kind leads to loss of muscle mass (Rolland 2008), while reduced physical activity is associated with an increase in sarcopenia measurements among elderly patients (Lee 2007, Viana 2015, Steffl 2017). Conversely, exercise can help reverse frailty indexes and maintain function in both humans (Reeves 2004, Koster 2012) and mice (Graber 2015, Madieros 2008).

Both aerobic and resistance exercise programs have generally been found to produce improvements in strength and function (Henderson 2017, Wanderley 2013, Villareal 2017, Yoo 2018). Some studies show aerobic exercise may result in greater all-around functional improvement (Wanderley 2013) and in particular improve mitochondrial-function-related factors (Yoo 2018), while resistance exercise produces greater improvements in muscle mass and strength (Landi 2014, Villareal 2017, Yoo 2018) as well as increases in bone density (Villareal 2013, Armamenta-Villareal 2019). Recently, a focus on power training or HIFT (high intensity functional training) which combine elements of aerobic and resistance training have shown these exercise modalities to potentially be superior in improving physical functional capacity (Graber 2019, 2. Ben-Zeev 2021, Feito,2018). However, the relationships between functional ability, age, and changes at the molecular level following exercise are complicated and not fully understood. In order to understand how to best maintain muscular function and maximize the number of years an individual may retain full functional ability; pre-clinical models are needed to determine direct cause and effect in mechanisms.

In this study we compare 6-10-month adult C57BL/6 mice (10m, the range correlating to mid-20’s to early 30s in a human) and 22-26-month-old elderly mice (26m, the range about 70-80-85 years in a human) to understand the effects of exercise on sarcopenia progression and decline of physical function. Mice underwent personalized endurance training via one of two types of aerobic exercise: voluntary wheel running (VWR, to simulate voluntary exercise) or treadmill running (HIIT, used to simulate high-intensity interval training). A previously published functional battery of tests (Graber 2015, 2020) is used to compile a composite functional score (CFAB, <u>c</u>omprehensive functional <u>a</u>ssessment <u>b</u>attery) to monitor how age and exercise regime affects functional abilities of mice. This data improves our understanding of the impact of aerobic exercise intensity on physical function, and notably that exercise type in older untrained mice seems to be less important to simply exercising itself for improving function, while mode does matter in adult mice.

## Methods

### Mice

Male C57BL/6 mice were obtained from the NIH NIA Aging Rodent Colony and Charles River Laboratory; and, were treated humanely under approved IACUC protocols. They were housed in groups, fed and watered *ad libitum*, and housed in 12-hour light/dark cycles at 22 °C. There were n=24 mice aged 6 months old and n=18 mice aged 22 months old at the start of the study. During the course of the study four of the younger adult mice and two of the older adult mice died of natural causes or were euthanized.

### Study Design

See **Figure 1** for study design schematic. After an appropriate acclimation period and application of identifying ear punches, mice spent an average of two weeks pre-testing for determination of functional abilities (see section on Functional Tests, CFAB, and Other Tests). Following pre-testing, mice were randomized into groups designated 10m (adult mice), 26m (older adult mice), RUN (voluntary wheel running), HIIT (high intensity interval training), and CON (sedentary control mice). See section on Exercise Training for more details. An exclusion criteria of >500 revolutions per week on the running wheels during pretesting was incorporated to ensure that any mice sent to the RUN group would actually use the wheel. Mice running below that threshold were automatically randomized into the HIIT or CON groups. After thirteen weeks of exercise, mice were post-tested for determination of functional abilities, then sacrificed for tissue collection and *in vitro* physiology.

**Figure 1.**
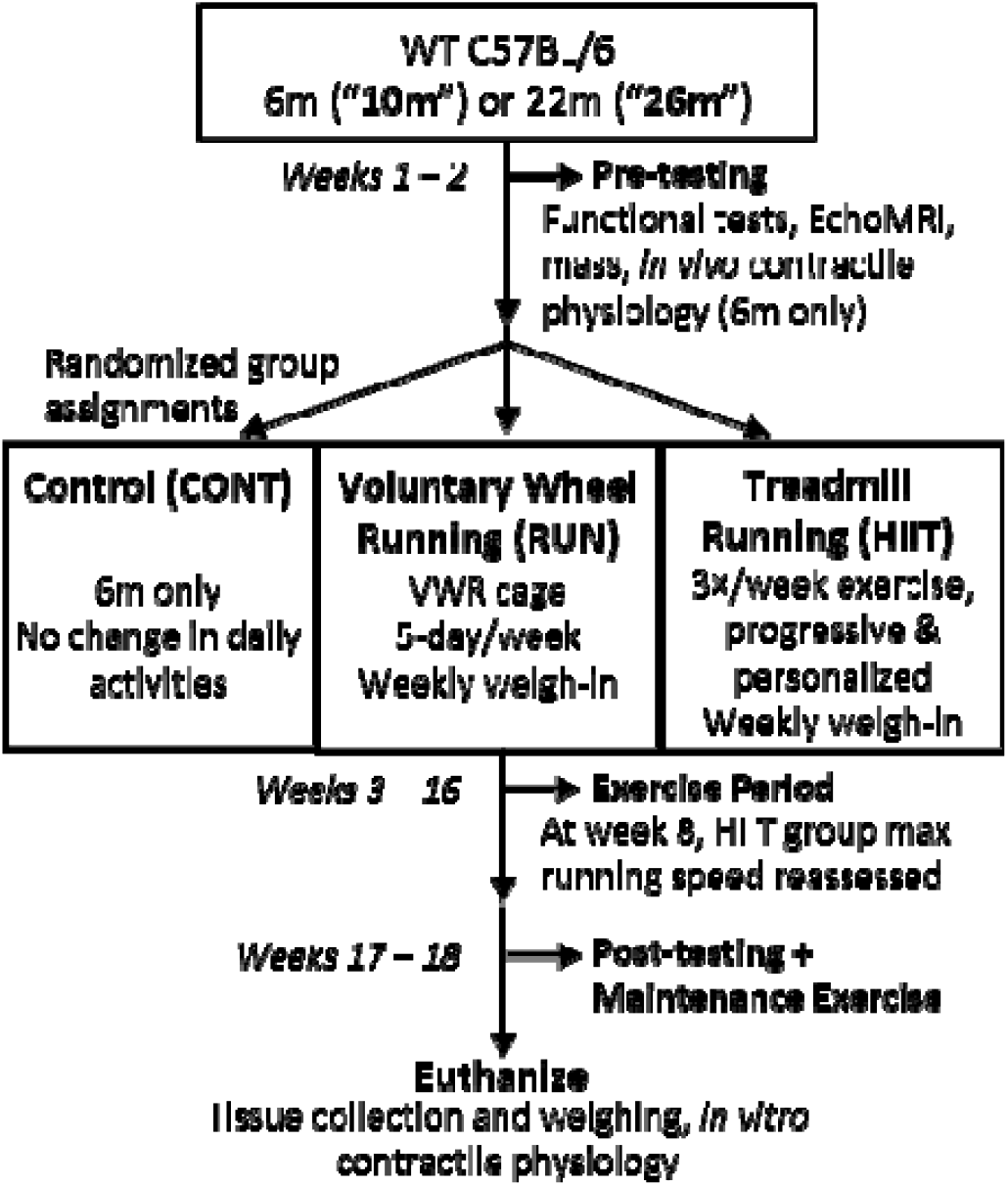
Study Design. Prior to exercise assignments, C57BL/6 mice aged 6 months (10m, n=24) or 22 months (26m, n=18) were pre-tested for functional abilities (voluntary wheel running, rotarod, treadmill, grip strength, inverted cling) and basic physiology (mass, body fat composition, *in vivo* physiology) for an approximate duration of two weeks. Afterwards, mice were randomized into volitional exercise/RUN (10m n=8, 26m n=8), treadmill running/HIIT (10m n=8, 26m n=10), or sedentary controls/CON (10m n=8). For the next thirteen weeks, RUN groups spent 5 days/week in cages equipped with wheels, HIIT groups spent 3 days/week undergoing progressive personalized treadmill running regimes, and CON groups lived in cages of up to three mice with no proscribed exercise. At the end of the exercise period, mice were post-tested for functional abilities and basic physiology before being euthanized for tissue collection and measurements of leg muscle mass. Because post-testing occurred over a period of time, maintenance exercise was provided for HIIT groups between the end of the exercise period and euthanization.

### Functional Tests

#### Neuromuscular Performance

Mouse functional ability and exercise capacity was assessed using the Comprehensive Functional Assessment Battery (CFAB) as previously described (Graber, 2020). Briefly, mice underwent a series of common well-validated functional tests (grip test, inverted cling, rotarod, voluntary wheel running, and treadmill running) at the beginning of the experiment (pre-training) and again at the conclusion (post-training). Each test score for individual mice was standardized (a difference in standard deviations from previously published six-month means) and summed to create a composite CFAB score as a reliable means of comparing physical function. Individual testing procedures are discussed briefly in the Online Supplement Methodology Section, and have been previously published (Graber, 2020; Graber, 2019; Graber, 2018; Graber, 2015; Graber, 2013).

### Other Tests

#### in vivo contractile physiology

We used an Aurora Whole Mouse 3-in-One physiology suite (Aurora Scientific). Methods have been published elsewhere (Graber 2019b; Graber 2020; Neelakantan 2019, Brightwell, 2021) and see further details in the online supplement. In brief, we anesthetized the mouse using isoflurane to remove conscious control of skeletal muscle. The mouse was placed on a heated platform, the foot set into a footplate attached to a force transducer, needle electrodes were placed so as to produce optimal twitch force in the plantar flexors (gastrocnemius complex), then optimal current was determined to produce the maximum twitch. We then made a force-frequency curve at a single pulse, 10 Hertz (Hz), 40 Hz, 80 Hz, 100 Hz, 120 Hz, 150 Hz, 180 Hz, 200 Hz, and a second single pulse to determine the maximum tetanic isometric torque (mN*m).

#### Body and Muscle Mass

Mass was measured during *in vivo* contractile physiology, at the time of tissue collection prior to euthanasia, and once a week during the treadmill and running wheel protocols. Additionally, an EchoMRI-700 (Echo Medical Systems) was used to determine body composition (fat percentage, fat%) at pre- and post-training.

### Exercise Training

The mice were randomly divided into groups: runners (RUN, both ages n=8, 7 survived to the study conclusion in the older mice and 6 in the adult mice), high intensity interval training (HIIT,10m, n=8; 26m, n=10, 9 survived to study end), and sedentary control (CON, 10m, n=8, 6 survived to study end). Sedentary mice were housed in cages with environmental enrichment but with no exercise proscribed. Exercise mice followed the below programs of exercise.

#### RUN (VWR)

VWR training protocols are the same as VWR functional testing, except that mice were housed in cages equipped with running wheels for five days, followed by a two-day period when mice were returned to social housing for rest and recuperation. Each RUN mouse completed one five-day session per week, with mass and total wheel revolutions recorded at the end of the five days (converted to km/day).

#### HIIT (treadmill running)

Using maximum running speed at failure from pre-training treadmill functional testing, HIIT mice were placed in similarly paced groups. The median max speed of each group was used to determine 75%, 80%, and 85% max speed for HIIT intervals. Mice trained three times per week with a two-day rest during weekends. HIIT sessions began with a one-minute 3 m/min warm-up walk, followed by intervals separated by 60 s walk steps (3 m/min). Each interval began with a 30 s acceleration step to running speed (set to 75%, 80%, or 85% max speed), which was maintained for 60 s, before ending with a 30 s deceleration back to walking speed. HIIT sessions ended with a two-minute 3 m/min cool-down walk. Initially, groups trained at lower-intensity, fewer HIIT intervals (three intervals at 75% max speed, or 75-75-75). Over time, interval number and running speed increased according to the abilities of the mice until groups were running 75-80-85-80-75 intervals. Halfway through the study, HIIT mice repeated the functional treadmill test and group speeds and assignments changed according to how the maximum running speed of each mouse had improved.

### Statistics and Data

Unless otherwise noted, the mean plus or minus the standard error of the mean (SEM) is used when reporting statistics and data. Statistical significance was set at p<0.05, and trends are reported where 0.05<p<0.10. ANCOVA (adjusted for body mass), ANOVA, or Student’s T-Test are used to compare means between subjects as appropriate. We used repeated measures ANCOVA (adjusted for body mass if appropriate), ANOVA, and paired t-tests to test with-in subject changes. We used least significant differences post-hoc testing. Some tests had outliers (defined as greater than >2 standard deviations from the mean, in some measurements much higher than 2), we analyzed both with and with these outliers and report both if needed for clarity. SPSS v27(IBM) were used to perform statistical analysis.

## Results

### Functional Tests

Pre-training baseline values and post-training functional testing value means of the mice are shown in **Table 1**, along with the main effects and interactions between terms when the data is analyzed as repeated measure ANOVA/ANCOVA (2×2×2: 2 ages, 2 exercises, pre- to post-training scores, the 10m CON group is excluded). See **Table S1** in the Supplementary Online Results for more details on specific values for mean percent change, and mean differences pre- to post-training, along with p-values. Herein we report the percent change from pre- to post-training (p-values are within group changes from 3×2×2 repeated measures ANCOVA, adjusted for pre- and/or post-training body mass if necessary, or ANOVA, post-hoc testing by repeated measures t-test). **Figure 2** depicts the mean differences from pre- to post-training and uses a 1-way univariate ANOVA, with the 10m sedentary control as the baseline, for statistical analysis.

**Table 1.**
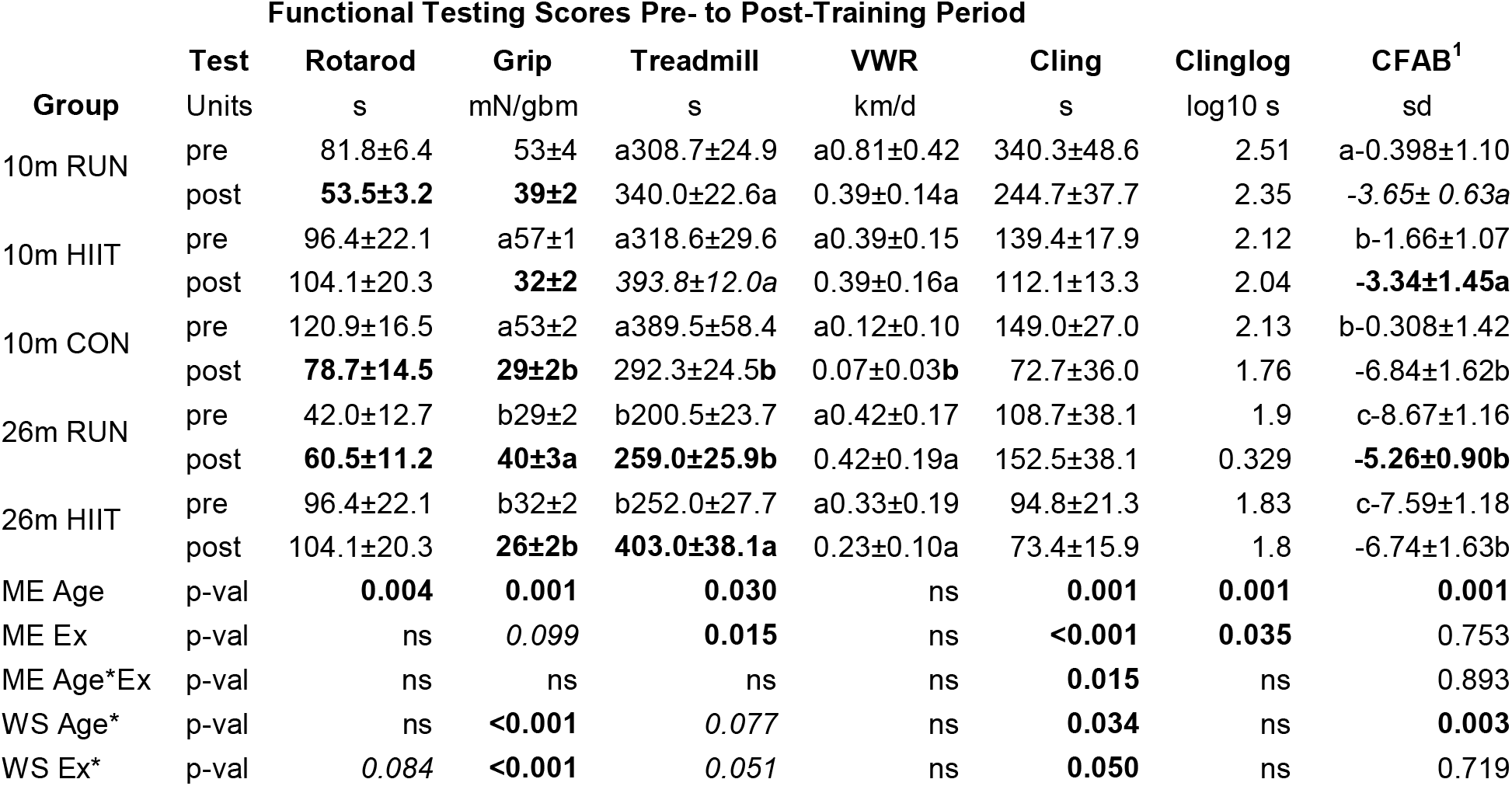
Functional Testing Score. Mean and standard error at pre-testing (pre) and post-training (post), statistics from paired t-test post hoc from Repeated Measures ANOVA (CFAB, Tread) or ANCOVA (others, adjusted for body mass). **Bold** indicates significance change (p<0.05) and *italics* indicates a trend (0.05<p<0.10), between groups; different letters indicate significant difference from Univariate ANOVA (CFAB, Tread) or ANCOVA (others, adjusted for body mass) with letters before comparing pre-tests and after comparing post-tests. ME = main effect between subjects, WS = main effect within subjects, all ME and WS from 2×2×2 Repeated Measures ANCOVA (adjusted for body mass) or ANOVA for CFAB (and tread), CON excluded.

| Functional Testing Scores Pre- to Post-Training Period |  |  |  |  |  |  |  |  |
| --- | --- | --- | --- | --- | --- | --- | --- | --- |
| Group | Test | Rotarod | Grip | Treadmill | VWR | Cling | Clinglog | CFAB<sup>1</sup> |
|  | Units | s | mN/gbm | s | km/d | s | log10 s | sd |
| 10m RUN | pre | 81.8±6.4 | 53±4 | a308.7±24.9 | a0.81±0.42 | 340.3±48.6 | 2.51 | a-0.398±1.10 |
|  | post | **53.5±3.2** | **39±2** | 340.0±22.6a | 0.39±0.14a | 244.7±37.7 | 2.35 | -3.65± 0.63a |
| 10m HIIT | pre | 96.4±22.1 | a57±1 | a318.6±29.6 | a0.39±0.15 | 139.4±17.9 | 2.12 | b-1.66±1.07 |
|  | post | 104.1±20.3 | **32±2** | *393.8±12.0a* | 0.39±0.16a | 112.1±13.3 | 2.04 | **-3.34±1.45a** |
| 10m CON | pre | 120.9±16.5 | a53±2 | a389.5±58.4 | a0.12±0.10 | 149.0±27.0 | 2.13 | b-0.308±1.42 |
|  | post | **78.7±14.5** | **29±2b** | 292.3±24.5b | 0.07±0.03b | 72.7±36.0 | 1.76 | -6.84±1.62b |
| 26m RUN | pre | 42.0±12.7 | b29±2 | b200.5±23.7 | a0.42±0.17 | 108.7±38.1 | 1.9 | c-8.67±1.16 |
|  | post | **60.5±11.2** | **40±3a** | **259.0±25.9b** | 0.42±0.19a | 152.5±38.1 | 0.329 | **-5.26±0.90b** |
| 26m HIIT | pre | 96.4±22.1 | b32±2 | b252.0±27.7 | a0.33±0.19 | 94.8±21.3 | 1.83 | c-7.59±1.18 |
|  | post | 104.1±20.3 | **26±2b** | **403.0±38.1a** | 0.23±0.10a | 73.4±15.9 | 1.8 | -6.74±1.63b |
| ME Age | p-val | **0.004** | **0.001** | **0.030** | ns | **0.001** | **0.001** | **0.001** |
| ME Ex | p-val | ns | 0.099 | **0.015** | ns | **<0.001** | **0.035** | 0.753 |
| ME Age*Ex | p-val | ns | ns | ns | ns | **0.015** | ns | 0.893 |
| WS Age* | p-val | ns | **<0.001** | 0.077 | ns | **0.034** | ns | **0.003** |
| WS Ex* | p-val | 0.084 | **<0.001** | 0.051 | ns | **0.050** | ns | 0.719 |

**Figure 2.**
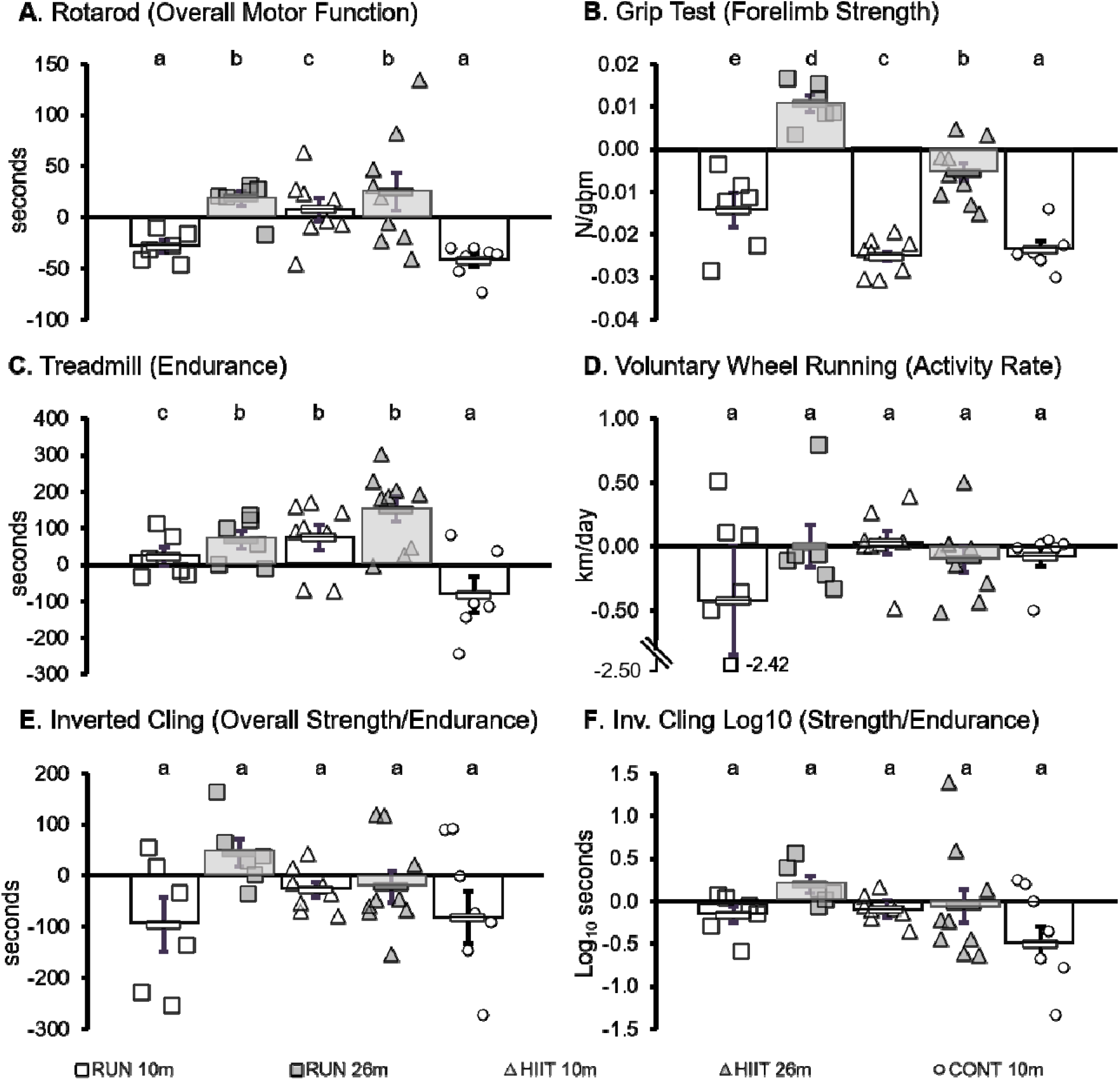
Functional Test Results. Difference in mouse performance for each functional test between pre-training and post-training. A) Rotarod functional test, data measured in seconds (s). B) Grip strength functional test, data measured in Newtons and reported after correction for body mass as Newtons per gram body mass (N/gbm). C) Treadmill functional test, data measured in seconds (s). D) Voluntary wheel running functional test, data measured as kilometers per day (km/day). E) Inverted cling functional test, data measured in seconds (s). F) Inverted cling functional test, data measured in seconds and reported after transformation to log10seconds. 10m VWR, open squares (n=7); 26m VWR, shaded squares (n=7); 10m HIIT, open triangles (n=8); 26m HIIT, shaded triangles (n=10); 10m CON, open circles (n=6). Each symbol represents one individual mouse, while bars and error bars represent average and standard error of the mean for each group. Different letters represent statistically significant differences (p<0.05) from the 10m CON group, calculated using a one-way univariate ANOVA with least significant differences post-hoc testing.

### Rotarod

Function Improved, Maintained, or Loss Mitigated with HIIT and 26m RUN

The rotarod is designed to test overall motor function, combining elements of balance, coordination, gait speed, and power generation, with the outcome measures as the latency to fall in seconds (s). 10m RUN decreased significantly by 35% (p=0.004). 26m RUN increased significantly by 44% (p=0.049). 10m HIIT increased numerically by 8% (p=0.520). 26m HIIT increased numerically by 39% (p=0.222). 10m CON decreased significantly by 37% (p<0.001).

### Grip Test

Strength Improved in 26m RUN, Loss Mitigated in 26m HIIT and 10m RUN

The grip test measures forelimb strength and is measured in Newtons (N, or milliNewtons, mN). To correct for effects of body mass (larger mice have stronger grip), max grip strength was divided by body mass at time of test (mN/gbm, grams body mass). Within the groups, from pre-training to post-training (from paired t-test): the 10m RUN decreased by 26% (p=0.013), 26m RUN increased by 38% (p=0.003),10m HIIT decreased by 44% (p<0.001), 26m HIIT decreased by 19% (p=0.046), and 10m CON decreased by 45% (p<0.001).

### Treadmill

Endurance Maintained or Improved with Exercise

The treadmill tests endurance capacity and is measured in seconds. Within the groups: 10m RUN increased numerically by 10% (p=0.390), 26m RUN increased significantly by 29% (p=0.043), 10m HIIT tended to increase by 44% (p<=0.063), 26m HIIT increased significantly by 60% (p=0.002), and 10m CON decreased numerically by 21% (p=0.156).

### Voluntary Wheel Running

Activity Maintained in All Groups

The voluntary wheel running test, or VWR, measures volitional exercise capacity and is measured in kilometers per day (km/d). There were no statistically significant changes to VWR within groups: 10m RUN decreased numerically by 52% (p=0.364), 26m RUN did not change (p=0.994), 10m HIIT increased numerically by 9% (p=0.724), 26m HIIT decreased numerically by 31% (, p=0.341), and 10m CON decreased numerically by 46% (p=0.334). There were no significant differences in pre-testing VWR performance within either age group, however in post-testing the 10m CON group was significantly 85% lower than the 10m RUN (p=0.041).

### Inverted Cling

Strength/Endurance Improved in 26 RUN

The inverted cling test measures strength in all four limbs, as well as endurance, and is measured in seconds. Within groups, 10m RUN decreased numerically by 28% (p=0.131), 26m RUN increased numerically by 40% (p=0.174), 10m HIIT decreased numerically by 20% (p=0.108), 26m HIIT decreased numerically by 23% (p=0.501), and 10m CON decreased numerically by 51% (p=0.287). In post-testing between groups 26m Run increased significantly compared to the other groups (p<0.001).

There were no significant differences in pre-testing grip performance within 26m, though the 10m RUN cling time mean was 154% and 128% significantly greater than the 10m HIIT and CON, respectively, in pre-testing (t-tests: p=0.001 and p=0.043).

### CFAB

Function was Improved, Maintained or Loss Mitigated with Exercise

CFAB is the composite score given from the sum of standardized scores from the rotarod (s), inverted cling (log10 s), grip test (N), treadmill (s), and voluntary running wheel (km/d). Mean differences from pre- to post-training are shown in **Figure 3**, with letters representing significant differences from a 1-way univariate ANOVA, using the 10m sedentary control as baseline. Within groups,10m RUN tended to decrease by 336% p=0.061), 26m RUN increased significantly by 341% (p=0.016), 10m HIIT decreased significantly by 279% (p=0.028), 26m HIIT increased numerically by 11% (p=0.657), and 10m CON decreased significantly by **2120%** (p=0.007), losing 6.53 sd over the course of the study.

**Figure 3.**
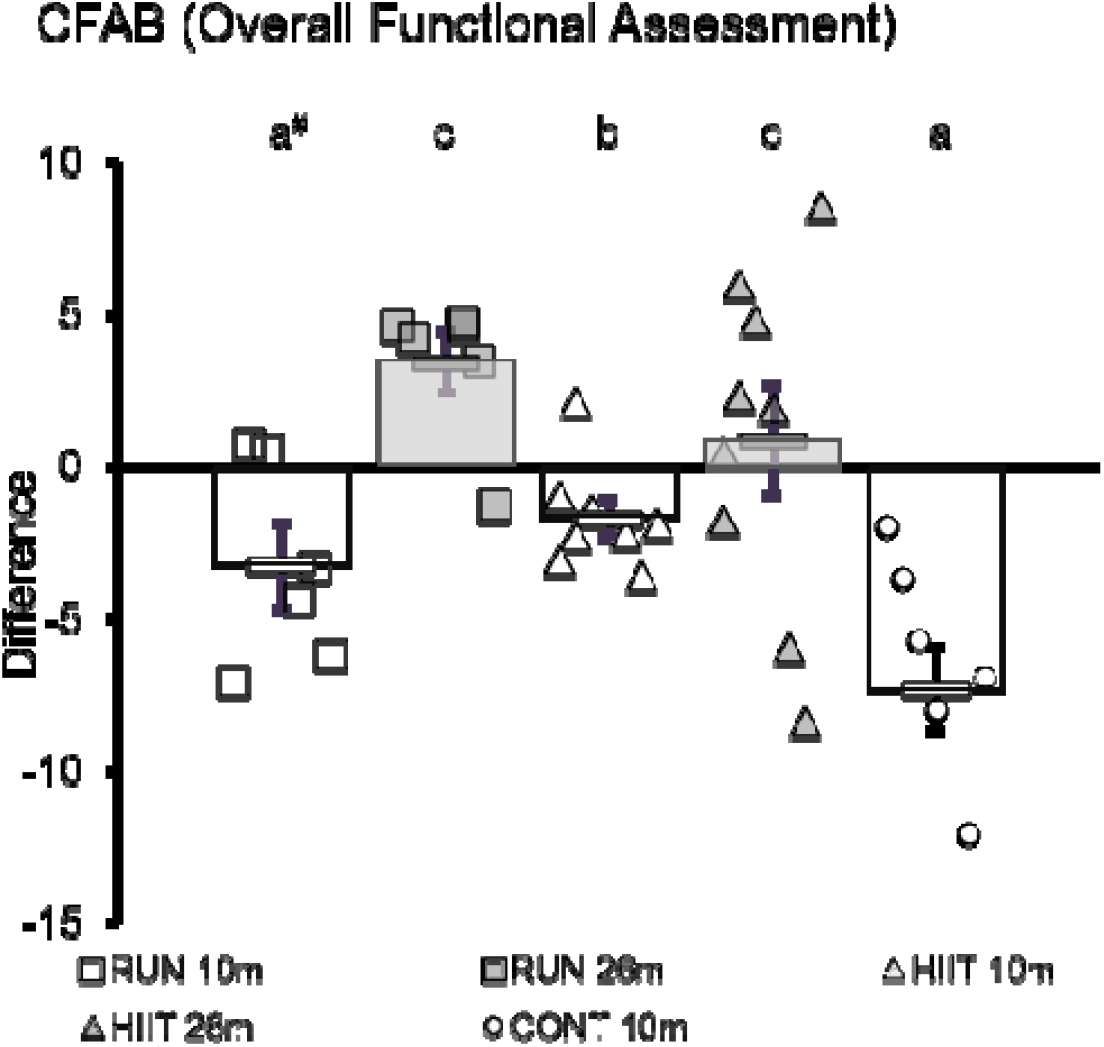
Comprehensive Functional Assessment Battery Results. Difference in CFAB between pre- and post-training. Data from functional tests were standardized to a previously published 6-month mean, then summed to calculate CFAB. A positive difference in CFAB indicates increased overall physical function between pre- and post-training; a negative difference indicates decreased overall physical function. 10m RUN, open squares (n=7); 26m RUN, shaded squares (n=7); 10m HIIT, open triangles (n=8); 26m HIIT, shaded triangles (n=10); 10m CON, open circles (n=6). Each symbol represents one individual mouse, while bars and error bars represent average and standard error of the mean for each group. Different letters represent statistically significant differences (p<0.05) from the 10m CON group, calculated using a one-way univariate ANOVA with least significant differences post-hoc testing.

The interactions of age*CFAB (p=0.034) and exercise type*age*CFAB (p=0.050) subjects, excluding CON from analysis (2×2×2 ANOVA), were significant, and between subjects both the main effects of age (p=0.001) and exercise (p<0.001), and the age*exercise interaction (p=0.015) were significant. There were no significant differences in pre-testing CFAB within 26m, though the 10m RUN CFAB mean was 154% and 128% significantly greater than the 10m HIIT and CON, respectively, in pre-testing (t-tests: p=0.001 and p=0.043).

### Body Composition

#### Fat %

Exercise Improved 26m Body Composition Improved; Mitigated 10m Increase in % Body Fat

Body composition altered markedly between the start and end of the experiment for all groups. P-values are from paired t-tests of the pre- to post-measurements. As shown in **Figure 4a**, 26m RUN mice lost an average of 13.7 ± 2.1% body fat (p=0.001, pre- to post-training), while 26m HIIT mice lost 6.2 ± 1.0% body fat (25.8% decrease, p<0.001). 10m groups generally gained body fat, with 10m RUN gaining 9.0 ± 2.1% fat (51.7% increase, p=0.008) and the 10m HIIT group gaining 12.1 ± 1.9% fat (100.9% increase, p<0.001). However, the 10m CON group gained 18.4 ± 2.1% body fat, a 160.5% increase (p<0.001), indicating that exercise mitigated increases in body fat associated with 6- to 10-month growth and development. See **Table S1.3** in the Supplement for raw data values.

**Figure 4.**
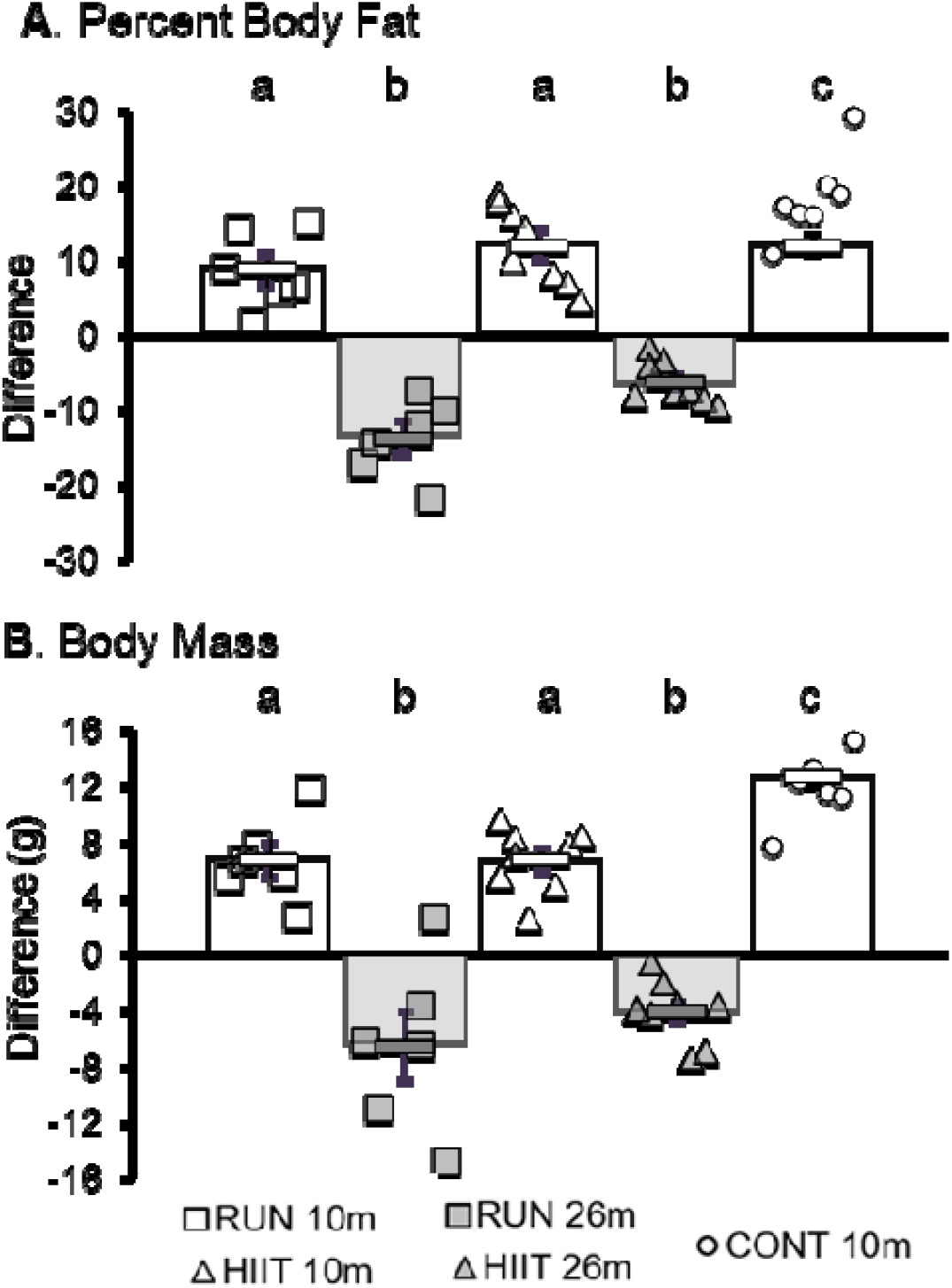
Body Composition. Changes in body composition between pre- and post-training. A) Difference in percent body fat, measured in grams using EchoMRI and reported after transformation to percent fat over total fat + lean mass. B) Difference in body mass, measured in grams. 10m RUN, open squares (n=7); 26m RUN, shaded squares (n=7); 10m HIIT, open triangles (n=8); 26m HIIT, shaded triangles (n=10); 10m CON, open circles (n=6). Each symbol represents one individual mouse, while bars and error bars represent average and standard error of the mean for each group. Different letters represent statistically significant differences calculated using a one-way univariate ANOVA with least significant differences post-hoc testing.

### Body Mass

Total body mass also changed between pre- and post-training (**Figure 4b**), particularly for 26m groups. The 26m RUN group lost an average of 7.0 ± 2.6 g of body mass (p=0.004) while the 26m HIIT group lost 4.8 ± 0.7 g (p,0.001), decrease of 14.8% and 12.1%, respectively. 10m groups body masses increased over the course of the experiment, with the RUN group increasing 6.6 ± 1.4 g (21.9% increase, p=0.003), the 10m HIIT group gaining 6.6 ± 0.7 g (21.7%, p<0.001), and the 10m CON gaining 11.0 ± 1.2 g (34.2%, p<0.001). See **Table S1.3** in the Supplement for raw data values.

### Muscle Mass

Hindlimb muscles were collected following euthanization and their masses measured (**Table 2**). RUN groups had larger gastrocnemius, plantaris, tibialis anterior (TA), extensor digitorum longus (EDL) than their age-matched HIIT counterparts. 10m CON mice had heavier absolute mass of gastrocnemius and plantaris than 10m RUN or HIIT, but after normalization to body mass they had lighter means for both muscles. The 10m CON had larger TA and EDL muscles compared to the 10m HIIT group, but these muscles were lighter compared to the 10m RUN group. When corrected for body mass, 10m CON mice had less massive soleus muscles than the other 10m groups. See **Figure 5** for details and statistics from One-Way Univariate ANOVAs.

**Table 2.** Muscle Mass Raw Data. Means ± standard error are shown. **Key:** gbm = grams of body mass, GAS = gastrocnemius, Plant = plantaris, TA = tibialis anterior, EDL = extensor digitorum longus, SOL = soleus, N. = normalized to body mass, TLM = total leg muscle mass (sum of GAS, PLANT, SOL, TA, and EDL).

| Muscle | Unit | 10m RUN | 26m RUN | 10m HIIT | 26m HIIT | 10m CON |
| --- | --- | --- | --- | --- | --- | --- |
| GBM | mg | 36.78 ± 1.93 | 35.93 ± 1.63 | 36.85 ± 1.37 | 34.85 ± 1.37 | 42.82 ± 1.80 |
| GAS | mg | 171.87 ± 4.51 | 143.79 ± 3.23 | 171.47 ± 5.40 | 131.02 ± 3.96 | 186.12 ± 6.13 |
| GAS N. | mg/gbm | 4.71 ± 0.15 | 4.03 ± 0.16 | 4.69 ± 0.18 | 3.81 ± 0.13 | 4.36 ± 0.14 |
| PLANT | mg | 21.65 ± 0.33 | 18.64 ± 0.61 | 21.39 ± 0.82 | 17.63 ± 0.77 | 22.95 ± 1.32 |
| PLANT N. | mg/gbm | 0.60 ± 0.03 | 0.52 ± 0.03 | 0.58 ± 0.02 | 0.51 ± 0.02 | 0.54 ± 0.02 |
| TA | mg | 51.83 ± 1.46 | 43.95 ± 3.36 | 48.05 ± 1.59 | 40.82 ± 1.77 | 51.28 ± 1.75 |
| TA N. | mg/gbm | 1.43 ± 0.10 | 1.22 ± 0.08 | 1.32 ± 0.07 | 1.19 ± 0.07 | 1.20 ± 0.02 |
| EDL | mg | 15.35 ± 3.13 | 12.27 ± 0.48 | 12.18 ± 0.52 | 11.21 ± 0.43 | 14.34 ± 1.29 |
| EDL N | mg/gbm | 0.43 ± 0.10 | 0.34 ± 0.01 | 0.33 ± 0.01 | 0.33 ± 0.02 | 0.34 ± 0.03 |
| SOL | mg | 12.90 ± 0.61 | 10.55 ± 0.75 | 11.43 ± 0.51 | 9.66 ± 0.53 | 11.91 ± 0.78 |
| SOL N. | mg/gbm | 0.35 ± 0.02 | 0.29 ± 0.02 | 0.31 ± 0.02 | 0.23 ± 0.02 | 0.28 ± 0.01 |
| TLM | mg | 529.03 ± 16.54 | 454.77 ± 12.74 | 527.87 ± 13.07 | 420.69 ± 12.83 | 570.88 ± 19.81 |
| TLM N. | mg/gbm | 14.49 ± 0.48 | 12.73 ± 0.40 | 14.44 ± 0.53 | 12.25 ± 0.46 | 13.37 ± 0.30 |

**Figure 5.**
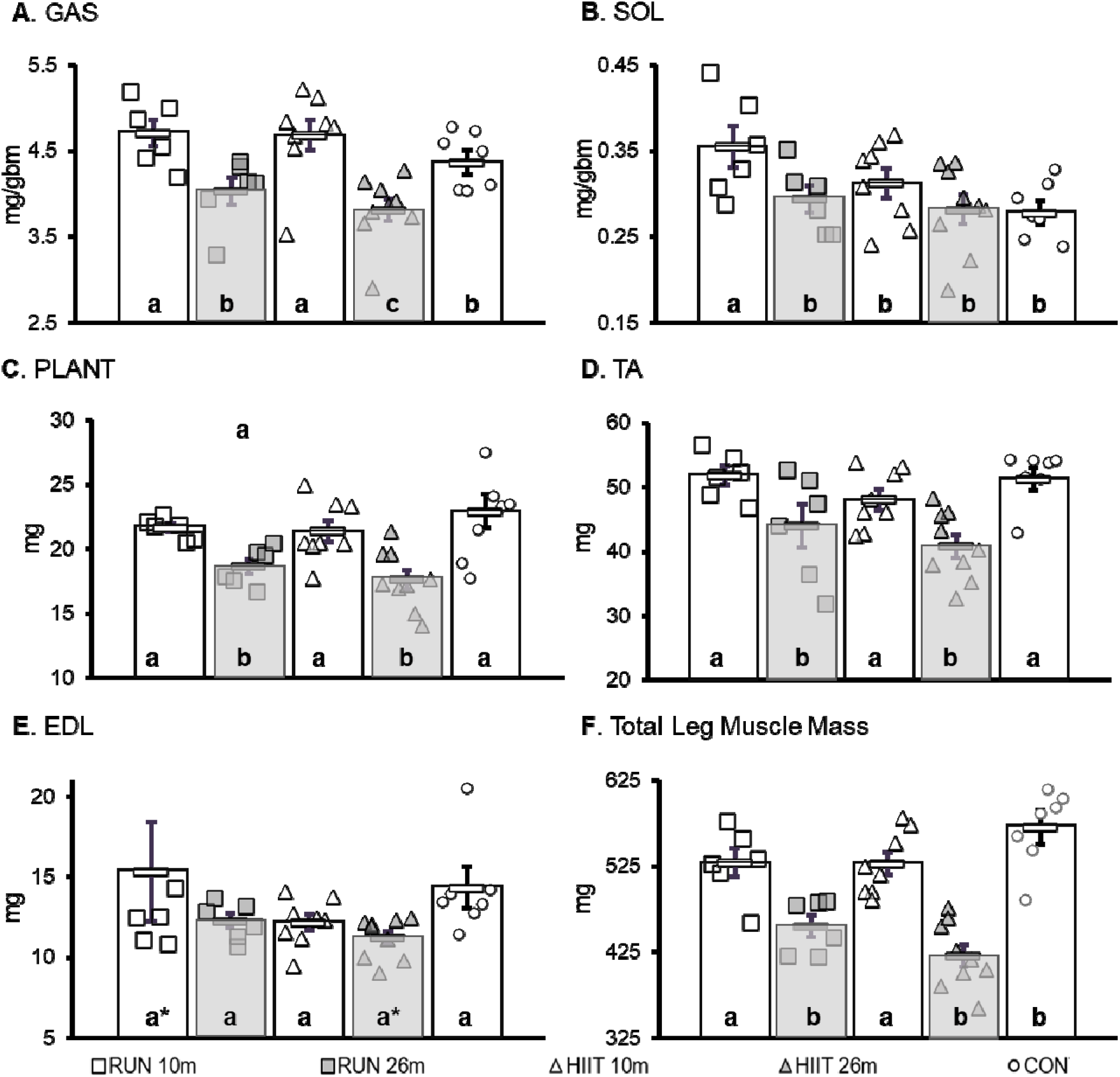
Muscle Mass. Postural muscles (SOL) and support of body weight during movement (GAS) were normalized to body mass (mg/gbm), all others are presented as absolute mass (mg). A) GAS (gastrocnemius). B) SOL (soleus). C) PLANT (Plantaris). (D)TA (tibialis anterior). E) EDL (extensor digitorum longus) (g). F) Mean total mass of leg muscles (g). **KEY:** mg = milligrams, gbm = grams of body mass, 10m RUN, open squares (n=7); 26m RUN, shaded squares (n=7); 10m HIIT, open triangles (n=8); 26m HIIT, shaded triangles (n=10); 10m CON, open circles (n=6). Each symbol represents one individual mouse, while bars and error bars represent average and standard error of the mean for each group. Different letters represent statistically significant differences calculated using a one-way univariate ANOVA with least significant differences post-hoc testing.

### Plantar Flexor Peak Tetanic Torque (*in vivo* Contractile Physiology)

See **Figure 6** for more details **and Table S1.3** in the Supplement for raw data values. NOTE: Because of technical limitations we do not have data for the older adult groups (26m). There was also a significant outlier in the CON group (−5.76% change from pre- to post, sd>3 from the mean) that increased variability and we report both with and without the outlier (**Figure 6** has the outlier designated, but the mean and statistics exclude it). Within the groups: From pre- to post-training all the 10m groups lost strength (normalized to body mass) with RUN decreasing by 0.057 ± 0.017 mN*m/g (−12.8%, p=0.043), HIIT decreasing by 0.089 ± mN*m/g (−23.8%., p=0.002), and without outlier CON decreasing by 0.118 ± 0.009 mN*m/g (−30.9%, p<0.003), with outlier CON decreased by 0.102 ± 0.018 mN*m/g (26.7%, p<0.001); statistics from paired t-tests of mean normalized tetanic force (pre- to post-training). With the outlier kept, there was no significant difference in the mean percent change (pre- to post-training) between groups using a one-way Univariate ANOVA (F= 2.605, p=0.105). However, without the outlier 10m RUN>10 HIIT>10 CON (F=4.704, p=0.026), with LSD posthoc testing 10m RUN>10m HIIT (RUN 11.0% difference, trend, p=0.064), 10m RUN > 10 CON (18.1% difference, p=0.009), 10m HIIT = 10 CON (numerical difference 7.0%, p=0.244).

**Figure 6.**
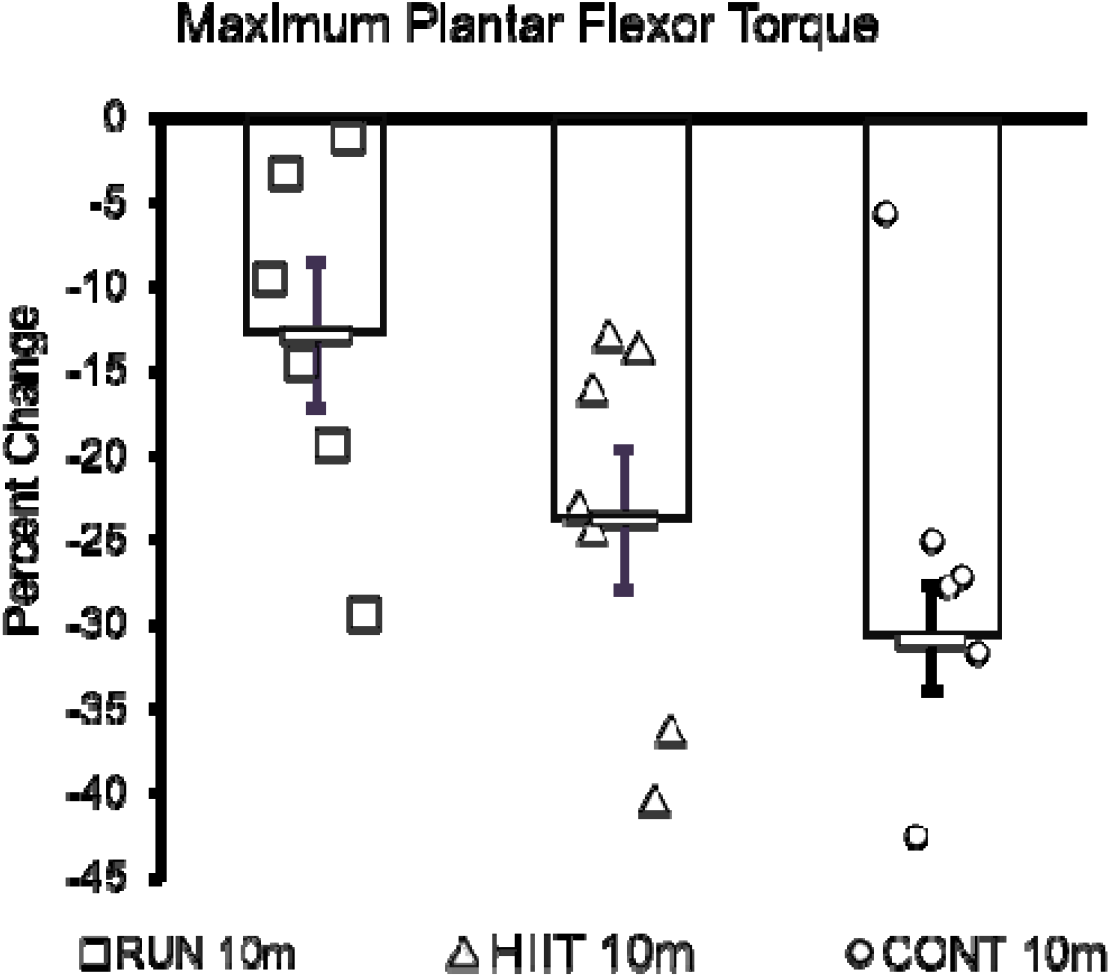
*in vivo* Contractile Physiology. Percent change in maximum plantar flexor torque between pre- and post-training (mean shown without the outlier in CON, -5.76% change >3 sd from the mean—marked with red circle). Mice were placed on heated platforms with one leg positioned with electrodes to produce maximum twitch force of the plantar flexors (gastrocnemius complex). Maximum tetanic isomeric torque was measured (mN*m) and is reported as percent change after correction to body mass (mN*m/gbm). Different letters represent statistically significant differences calculated using a one-way univariate ANOVA with least significant differences post-hoc testing (-outlier).

## Discussion

As we grow older, we begin to experience declining physical function. This is in part due to sarcopenia which is associated with loss of muscular strength, endurance, and mobility (McPhee 2016, Nascimento 2019). Declining physical function can lead to a lowered ability to perform activities of daily living and a reduced quality of life (McFee 2016, Marzetti 2017). Eventually, loss of function leads to loss of independence, requiring assisted living, such as placement into a skilled nursing facility for example (Marzetti 2017). Exercise, while not a cure, has shown promise as a potential therapy to reduce the rate of functional loss with age and improve quality of life.

Exercise has been shown to increase functional ability in older adults while also improving many biomarkers of physical fitness such as strength, muscle mass, VO_2max_ (Fiaterone 1990, Bonnefoy 2003 McPhee 2016). Valid pre-clinical models improve capacity to investigate the molecular mechanisms underlying age-associated functional loss and the age/exercise interaction. Therefore, in this study we designed aerobic exercise training programs for C57BL/6 mice to mimic high intensity interval training and voluntary activity (i.e. walking or jogging) often suggested as appropriate training programs for humans looking to improve or maintain physical performance. Using our CFAB scoring system (Graber 2020) we determined the effect of the two exercise types on functional capacity in both older adult and adult mice, using a sedentary adult control group to represent the archetypical “couch potato”. In the next sections we detail how our results compare to other findings in humans and rodents and cover differential outcomes by exercise type and age-group.

### Mouse Exercise

Our results support a growing base of knowledge that exercise improve measurements of health and function (Seldeen 2018, Graber 2015, Garcia-Valles 2013). Using a similar, though un-personalized high intensity interval training protocol, Seldeen et al. have previously shown that high intensity interval training in 24-month-old mice resulted in improved frailty scores (Seldeen 2018), just as we observed. Additionally, like our mice, they found that after sixteen weeks of HIIT training, aged mice spent significantly more time on the treadmill after HIIT training. Voluntary wheel running in mice has previously been shown to improve body composition (Manzaneres 2019), as we observed. Our observations of improved physical function following voluntary wheel running exercise are like those previously published (Graber 2015) for similarly aged mouse groups. Unlike our findings, the “old” exercise group in that study exhibited a decrease in frailty index; however, it should be noted that those mice were older than our group, and exercise only lasted four weeks. Thus, our recent results are more indicative of the effects of habitual exercise on a slightly younger (though still aged) group.

### Human Exercise

Proscribed high intensity exercise has been previously shown to improve measures of frailty in elderly humans, including improvements in the strength and flexibility (Hess 2006, Buckinx 2018), stair climbing (Bonnefoy 2003, Melzer 2009), walking time (Bonnefoy 2003), chair rising (Bonnefoy 2003, Buckinx 2018), and hand grip strength (Jimenez-Garcia 2018). Additionally, high intensity exercise in humans has long been shown to improve a number of markers of frailty and sarcopenia, including oxygen consumption, cardiovascular function, blood pressure, and inflammation (Buckinx 2018, Fiaterone 1990, Bonnefoy 2003, McPhee 2016). Voluntary wheel running exercise in mice is more complex to compare to human studies; however, clinical studies incorporating self-reported activity levels indicate that, like the mice in this study, more frequent daily activity is associated with improved frailty scores (Manini 2012, Tak 2013). Thus, one might think of voluntary wheel running as translatable to a study on older adults where the subjects are told to start walking or jogging (and wear a pedometer to track effort), while leaving it up to them to determine intensity of effort and volume (Teixeiria 2021; Cheatham 2018) Altogether, these data show that our models of exercise are comparable to results found in humans, supporting the supposition that samples collected from this study will provide valuable insights into mechanisms underlying physiological responses to exercise.

### VWR vs. HIIT

Contrary to our hypothesis, HIIT groups did not universally appear to gain greater exercise benefits than RUN groups. 26m HIIT and 26m RUN mice showed similar improvements in endurance and overall motor function (**Figure 2C**) as well as CFAB improvement (**Figure 3**). The only functional test in which 26m RUN and 26m HIIT showed different levels of improvement was the grip test, where 26m RUN mice exhibited improved forelimb strength (**Figure 2B**), a surprising result which may be cause by the running wheels having small ridges that the mice might grip onto as they run. Additionally, both 26m groups showed similar improvements in body composition (**Figure 3**). This suggests that for older age groups, any type of physical activity will result in improved functional performance. Commonly, “any exercise is better than no exercise” is a mantra used to encourage exercise participation by sedentary older adults, and the current study has demonstrated this in the context of older mice.

For the 10m RUN and HIIT groups, the 10m HIIT group performed better on the rotarod (Figure 2A) and treadmill tests (Figure 2C) at post-testing than the 10m RUN group. Additionally, while both 10m groups showed a decrease in CFAB from pre- to post-testing, the 10m HIIT group decreased significantly less. Like the 26m groups, the 10m RUN group had significantly stronger forelimbs than the 10m HIIT group (Figure 2B). All 10m groups showed reduced maximum plantar flexor torque post-training; however, the 10m RUN group decreased less than either the 10m HIIT or CON groups. Overall, this suggests that, excepting forelimb strength and maximum torque, high-intensity interval training did work better than voluntary running to mitigate loss of functional ability as mice aged from 6 months to 10 months. It is notable that exercise type did not affect changes in body composition for 10m groups compared to the 10m CON, other than to reduce overall change in body mass (CON gained both fat mass and muscle mass). Thus, exercise type may be more influential in less detrained adult mice, though we were highly conservative in the progressive difficulty during the HIIT protocol in the older mice to prevent any chance of injury and may not have challenged them sufficiently. We had a similar issue in previous work on power training (Graber, 2019a), thus in the future we may need to be more aggressive in increasing intensity and volume for older exercise groups.

### Older Adult Mice versus Adult Mice

It is apparent from our data that the effects of exercise depend on the age of the exercising group (exercise * age interaction. 10m groups generally showed greater benefits from high-intensity interval training. All 10m groups exhibited worse performance on functional tests at post-training, likely due to physiological transitions from 6 months of age (equivalent to a young adult) to 10 months of age (early middle-age). However, this decrease in performance and CFAB score showed greater mitigation from HIIT exercise. Exercise type did not significantly affect 10m body composition. However, the older 26m group demosntrated a very different benefit profile. First, either voluntary running or high-intensity interval training had the same overall effect on functional performance and CFAB. Second, exercise did have a significant effect on body composition, resulting in decreased body fat percent and increased lean body mass. Third, while 10m groups experienced a decrease in functional performance between pre- and post-testing, 26m groups showed improvements in performance. This is likely due to the fact that prior to 22 months of age, the 26m groups were sedentary, leaving them with more room for improvement compared to the 10m groups at their 6-month prime-of-life point. It would be enlightening to follow an exercise group during a longitudinal study to observe how a lifetime of exercise affects functional abilities at a similar post-training testing point.

### Caveats

One caveat that we must address is that due to complications related to the SARS-CoV-2 global pandemic and the ensuing shut-downs, we were unable to supervise a 26m CON group or perform in vivo contractile on the 26m exercise groups. Thus, we do not know what change would have occurred in a 26m CON group over the 4-month period. We do know from this study that the in the 10m CON we saw a large degree of degradation in functional capacity that was offset by exercise, and in the past we and others have shown that various forms of exercise do either improve function or mitigate the degree of functional loss versus sedentary controls in older adult mice (Graber, 2015; Graber, 2019a; Seldeen 2018; Seldeen 2019). Secondly, we may have erred on the side of caution in advancing the HIIT protocol intensity. In future work, a dose-response study is warranted to determine if greater intensity would increase gains.

### Future Directions

With these valid exercise models, future studies will be conducted to examine the interaction between functional loss, exercise and age at the level of the proteome and transcriptome to gain insight why some individuals respond better to exercise than others, and whether there is a difference in exercise response when older that limits gains (so-called anabolic resistance). Using samples collected from the mice in this study, we will look to understand the molecular mechanisms associated with declining function, determine which ones are mitigated by exercise, and seek to uncover potential novel therapeutic targets.

## Conclusion

As the world population ages, it will become increasingly necessary to find ways to maintain physical function and independence for as long as possible. Exercise is an important component of this, but more information is needed to understand exactly how exercise affects sarcopenia and frailty, and eventually to find ways to mimic these effects in individuals who are unable to participate in a standard exercise program. Here, we have shown that exercise improves overall physical function in both adults and older adults, but the pattern of improvement differs according to age. Younger adults benefit more from high-intensity interval training while older adults receive similar functional benefits from high-intensity exercise and lower-intensity voluntary exercise. This data will be compared to studies of the molecular mechanisms present in samples from these mice, allowing us to make direct comparisons between function, exercise, and key pathways in muscular function and development.

## Acknowledgments

We would like to acknowledge Hill Hollaway and Evin Flinchum for technical assistance.

## Author Contributions

Conceptualization: T.G.G.; methodology: T.G.G., M.P.; validation: T.G.G., formal analysis: T.G.G., C.B., M.P.; investigation: T.G.G., M.P., C.B. E.S., A.S., A.F., N.N.; resources: T.G.G.; writing—original draft: M.P., C.B., A.S., E.S., A.F., N.N., T.G.G.; writing—review and editing: T.G.G., M.P., C.B., A.S., E.S., N.N., and A.F.; supervision: T.G.G., M.P.; project administration: T.G.G., M.P.; funding acquisition: T.G.G.

## Table Legends

**Table 1 Functional Testing Score.** Mean and standard error at pre-testing (pre) and post-training (post), statistics from paired t-test post hoc from Repeated Measures ANOVA (CFAB) or ANCOVA (others, adjusted for body mass). **Bold** indicates significance change (p<0.05) and *italics* indicates a trend (0.05<p<0.10). ME = main effect, all ME from 3×2 Repeated Measures ANCOVA (adjusted for body mass) or ANCOVA for CFAB.

