## Supplemental Files for "Endurance Exercise to Improve Physical Function in Adult and Older Mice: High Intensity Interval Training (HIIT) versus Voluntary Wheel Running (VWR)"

**Table of Contents** **Pages**

| | |
|---|---|
| **Methodology: Functional Testing** | **2** |
| **Table S1.1 Functional Testing Raw Data Part 1** | **3** |
| **Table S1.2 Functional Testing Raw Data Part 2** | **4** |
| **Table S1.3 Body Composition and Plantar Flexor Torque Raw Data** | **5** |
| **References** | **6** |

### Online Only Supplemental Methodology:

**Functional Testing** (etailed protocols have been previously published at: Graber, 2020; Graber, 2019; Graber, 2018, Graber, 2015; and Graber, 2013)

*Rotarod (overall motor function):*

Mice were acclimated over two days to use a Panlab LE820 rotarod (Harvard Apparatus). On the third day, the rotarod was set to start at a speed of 4 rpm and accelerate over a total period of 300 s. Latency to fall was recorded for each mouse and the test repeated for a total of three trials, with a minimum fifteen-minute rest between tests. The longest latency period was used for CFAB score determination.

*Grip Test (forelimb strength):*

To measure forelimb strength, mice gripped a grip strength apparatus (BioSeb) set up with a grip bar. Mice were pulled straight backwards by their tail at a gentle, steady rate until they let go. Grip strength was recorded in newtons and the test was repeated for five total measurements. The highest grip force was used for CFAB score determination.

*Treadmill (endurance / volitional fatigue):*

Mice were trained over two days to use a treadmill (Columbus Instruments Exer 3/6) for a target minimum of two minutes per session. The treadmill shock grid was set to administer 3 Hz shocks at a repetition rate of 2 Hz, for a maximum of six shocks or three visits to the shock grid. On the third day, starting at a speed of 3 m/min and accelerating 0.6 m/min every 20 s, the time for each mouse to max out shocks was recorded, as well as the speed at shock-out. The time to shock-out was used for CFAB determination.

*Inverted Cling (overall strength / endurance):*

To test four-limb strength and endurance, mice were placed on a grid suspended above a padded floor. Latency to fall was recorded. After a minimum fifteen-minute rest, the test repeated, and the longer latency period used for CFAB score determination.

*Voluntary Wheel Running (activity and volitional exercise rate):*

Each mouse was placed in a cage equipped with a running wheel (Columbus Instruments) for seven days. Each running wheel was attached to a magnetic revolution counter. At the end of the seven days, total wheel turns were recorded and converted to km/day for CFAB score determination.

| **CFAB, sd** | **Pre-test** | **sd** | **SEM** | **Post-test** | **sd** | **SEM** | **Difference** | **sd** | **SEM** | **Percent Change** |
|---|---|---|---|---|---|---|---|---|---|---|
| 10m RUN | -0.398 | 2.689 | 1.098 | -3.649 | 1.531 | 0.625 | -3.251 | 3.312 | 1.352 | n/a |
| 26m RUN | -9.244 | 2.620 | 0.926 | -5.256 | 2.198 | 0.897 | 3.418 | 2.342 | 0.956 | n/a |
| 10m HIIT | -1.664 | 3.023 | 1.069 | -3.365 | 4.086 | 1.445 | -1.700 | 1.742 | 0.616 | n/a |
| 26m HIIT | -7.438 | 3.364 | 1.064 | -6.740 | 4.890 | 1.630 | 0.845 | 5.501 | 1.834 | n/a |
| 10m CONT | -0.308 | 3.472 | 1.417 | -6.836 | 3.969 | 1.620 | -7.306 | 3.106 | 1.389 | n/a |
| **Rotarod, s** | **Pre-test** | **sd** | **SEM** | **Post-test** | **sd** | **SEM** | **Difference** | **sd** | **SEM** | **Percent Change** |
| 10m RUN | 81.8 | 15.7 | 6.4 | 53.5 | 7.7 | 3.2 | -28.3 | 14.1 | 5.8 | -33.2 |
| 26m RUN | 44.3 | 26.9 | 9.5 | 60.5 | 27.4 | 11.2 | 18.5 | 17.5 | 7.1 | 382.2 |
| 10m HIIT | 96.4 | 62.4 | 22.1 | 104.1 | 57.5 | 20.3 | 7.8 | 32.4 | 11.4 | 41.4 |
| 26m HIIT | 64.0 | 29.5 | 9.3 | 87.6 | 60.0 | 20.0 | 24.8 | 56.2 | 18.7 | 164.5 |
| 10m CONT | 120.9 | 43.6 | 16.5 | 78.7 | 38.3 | 14.5 | -42.1 | 16.1 | 6.1 | -37.1 |
| **Grip Strength, N** | **Pre-test** | **sd** | **SEM** | **Post-test** | **sd** | **SEM** | **Difference** | **sd** | **SEM** | **Percent Change** |
| 10m RUN | 0.053 | 0.010 | 0.004 | 0.039 | 0.005 | 0.002 | -0.014 | 0.009 | 0.004 | -25.5 |
| 26m RUN | 0.028 | 0.004 | 0.001 | 0.040 | 0.007 | 0.003 | 0.011 | 0.005 | 0.002 | 36.8 |
| 10m HIIT | 0.057 | 0.004 | 0.001 | 0.032 | 0.004 | 0.002 | -0.025 | 0.004 | 0.001 | -44.1 |
| 26m HIIT | 0.032 | 0.006 | 0.002 | 0.026 | 0.007 | 0.002 | -0.006 | 0.007 | 0.002 | -16.7 |
| 10m CONT | 0.053 | 0.005 | 0.002 | 0.029 | 0.006 | 0.002 | -0.024 | 0.005 | 0.002 | -44.5 |
| **Treadmill, s** | **Pre-test** | **sd** | **SEM** | **Post-test** | **sd** | **SEM** | **Difference** | **sd** | **SEM** | **Percent Change** |
| 10m RUN | 308.3 | 72.2 | 29.5 | 331.5 | 60.6 | 24.7 | 23.2 | 60.4 | 24.6 | 12.0 |
| 26m RUN | 181.9 | 67.0 | 23.7 | 268.8 | 69.5 | 28.4 | 68.3 | 62.3 | 25.4 | 44.7 |
| 10m HIIT | 318.6 | 83.8 | 29.6 | 393.8 | 33.9 | 12.0 | 75.1 | 96.3 | 34.0 | 31.2 |
| 26m HIIT | 255.7 | 79.2 | 25.1 | 403.0 | 114.3 | 38.1 | 151.0 | 103.7 | 34.6 | 70.2 |
| 10m CONT | 389.5 | 62.0 | 25.3 | 306.4 | 66.3 | 25.1 | -81.7 | 119.9 | 48.9 | -17.9 |

**Table S1.1 Functional Testing Raw Data Part 1.** Group means, standard deviations, and standard error of the means for CFAB, functional tests (rotarod, grip strength, VWR, treadmill, inverted cling, log10 inverted cling), body composition (total body mass, percent body fat), and maximum plantar flexor torque. Values are presented for pre-training, post-training, pre- to post-training difference, and pre- to post-training percent chang

| **VWR, km/day** | **Pre-test** | **sd** | **SEM** | **Post-test** | **sd** | **SEM** | **Difference** | **sd** | **SEM** | **Percent Change** |
|---|---|---|---|---|---|---|---|---|---|---|
| 10m RUN | 0.814 | 1.039 | 0.424 | 0.389 | 0.330 | 0.135 | -0.425 | 1.044 | 0.426 | 14.6 |
| 26m RUN | 0.464 | 0.372 | 0.131 | 0.419 | 0.468 | 0.191 | 0.001 | 0.401 | 0.164 | 53.5 |
| 10m HIIT | 0.363 | 0.748 | 0.264 | 0.396 | 0.801 | 0.283 | 0.033 | 0.254 | 0.090 | 5674.8 |
| 26m HIIT | 0.300 | 0.379 | 0.120 | 0.226 | 0.291 | 0.097 | -0.102 | 0.301 | 0.100 | 7.3 |
| 10m CONT | 0.120 | 0.261 | 0.099 | 0.065 | 0.076 | 0.031 | -0.073 | 0.209 | 0.085 | 1096.8 |
| **Inv. Cling, s** | **Pre-test** | **sd** | **SEM** | **Post-test** | **sd** | **SEM** | **Difference** | **sd** | **SEM** | **Percent Change** |
| 10m RUN | 340.3 | 119.0 | 48.6 | 244.7 | 92.2 | 37.7 | -95.7 | 129.8 | 53.0 | -21.6 |
| 26m RUN | 90.4 | 85.9 | 30.4 | 152.5 | 93.2 | 38.1 | 43.8 | 67.8 | 27.7 | 80.8 |
| 10m HIIT | 139.4 | 50.6 | 17.9 | 112.1 | 37.7 | 13.3 | -27.3 | 41.8 | 14.8 | -14.7 |
| 26m HIIT | 92.7 | 60.7 | 19.2 | 73.4 | 47.7 | 15.9 | -21.3 | 90.9 | 30.3 | 262.7 |
| 10m CONT | 149.0 | 71.6 | 27.0 | 72.7 | 88.1 | 36.0 | -82.2 | 124.4 | 50.8 | -42.7 |
| **Log10 Inv. Cling** | **Pre-test** | **sd** | **SEM** | **Post-test** | **sd** | **SEM** | **Difference** | **sd** | **SEM** | **Percent Change** |
| 10m RUN | 2.508 | 0.160 | 0.066 | 2.353 | 0.211 | 0.086 | -0.155 | 0.246 | 0.100 | -5.9 |
| 26m RUN | 1.809 | 0.371 | 0.131 | 2.094 | 0.330 | 0.135 | 0.199 | 0.239 | 0.098 | 12.2 |
| 10m HIIT | 2.123 | 0.141 | 0.050 | 2.029 | 0.144 | 0.051 | -0.094 | 0.157 | 0.056 | -4.2 |
| 26m HIIT | 1.831 | 0.452 | 0.143 | 1.773 | 0.315 | 0.105 | -0.054 | 0.670 | 0.223 | 13.0 |
| 10m CONT | 2.131 | 0.209 | 0.079 | 1.649 | 0.454 | 0.185 | -0.494 | 0.562 | 0.229 | -21.8 |

**Table S1.2 Functional Testing Raw Data Part 2.** Group means, standard deviations, and standard error of the means for CFAB, functional tests (rotarod, grip strength, VWR, treadmill, inverted cling, log10 inverted cling), body composition (total body mass, percent body fat), and maximum plantar flexor torque. Values are presented for pre-training, post-training, pre- to post-training difference, and pre- to post-training percent chan

| Body Mass, g | Pre-test | sd | SEM | Post-test | sd | SEM | Difference | sd | SEM | Percent Change |
|---|---|---|---|---|---|---|---|---|---|---|
| 10m RUN | 30.168 | 2.664 | 1.088 | 36.957 | 3.933 | 1.606 | 6.788 | 3.012 | 1.230 | 22.7 |
| 26m RUN | 43.684 | 7.354 | 2.600 | 36.453 | 4.653 | 1.900 | -6.518 | 5.968 | 2.436 | -13.7 |
| 10m HIIT | 30.256 | 2.403 | 0.850 | 37.063 | 4.224 | 1.493 | 6.806 | 2.306 | 0.823 | 22.3 |
| 26m HIIT | 39.548 | 6.768 | 2.140 | 35.716 | 7.326 | 2.442 | -4.017 | 2.123 | 0.708 | -10.3 |
| 10m CONT | 31.979 | 1.421 | 0.537 | 44.285 | 2.986 | 1.219 | 12.748 | 1.656 | 0.741 | 39.4 |
| **Body Fat, %** | **Pre-test** | **sd** | **SEM** | **Post-test** | **sd** | **SEM** | **Difference** | **sd** | **SEM** | **Percent Change** |
| 10m RUN | 19.732 | 5.876 | 2.399 | 28.705 | 4.738 | 1.934 | 8.973 | 5.150 | 2.102 | 51.7 |
| 26m RUN | 32.777 | 7.107 | 2.513 | 18.709 | 6.708 | 2.739 | -13.736 | 5.201 | 2.123 | -42.4 |
| 10m HIIT | 15.109 | 6.630 | 2.344 | 27.244 | 7.425 | 2.625 | 12.135 | 5.384 | 1.904 | 101.0 |
| 26m HIIT | 25.858 | 8.258 | 2.753 | 20.375 | 9.545 | 3.182 | -6.243 | 2.877 | 1.017 | -25.8 |
| 10m CONT | 13.980 | 5.293 | 2.000 | 32.377 | 4.348 | 1.644 | 18.397 | 5.608 | 2.120 | 160.5 |
| **Max Plantar Flexor Torque, mN*m/g** | **Pre-test** | **sd** | **SEM** | **Post-test** | **sd** | **SEM** | **Difference** | **sd** | **SEM** | **Percent Change** |
| 10m RUN | 0.368 | 0.047 | 0.019 | 0.321 | 0.060 | 0.025 | -0.047 | 0.042 | 0.017 | -12.8 |
| 26m RUN | n/a | | | | | | | | | |
| 10m HIIT | 0.382 | 0.041 | 0.015 | 0.282 | 0.042 | 0.016 | -0.089 | 0.044 | 0.017 | -23.8 |
| 26m HIIT | n/a | | | | | | | | | |
| 10m CONT | 0.383 | 0.027 | 0.010 | 0.267 | 0.039 | 0.017 | -0.118 | 0.021 | 0.009 | -30.9 |

**Table S1.3 Body Composition and Plantar Flexor Torque Raw Data**. Group means, standard deviations, and standard error of the means for CFAB, functional tests (rotarod, grip strength, VWR, treadmill, inverted cling, log10 inverted cling), body composition (total body mass, percent body fat), and maximum plantar flexor torque. Values are presented for pre-training, post-training, pre- to post-training difference, and pre- to post-training percent change.

### Online Only Supplemental References:

1. Graber TG, Ferguson-Stegall L, Liu H, Thompson LV. Voluntary aerobic exercise reverses frailty in old mice. J Gerontol A Biol Sci Med Sci. 2015;70(9):1045–1058. doi:10.1093/gerona/glu163
2. Graber TG, Ferguson-Stegall L, Kim J-H, Thompson LV. C57BL/6 Neuromuscular Healthspan Scoring System. The Journals of Gerontology. Series A, Biological Sciences. 68(11): 1326-1336. doi: 10.1093/gerona/glt032
3. Graber TG, Fry CS, Brightwell CR, Moro T, Maroto R, Bhattarai N, Porter C, Wakamiya M, Rasmussen BB. Skeletal Muscle Specific Knockout of DEP domain-containing 5 Increases mTORC1 Signaling, Muscle Cell Hypertrophy, and Mitochondrial Respiration. J Biol Chem. 2019 Mar 15;294(11):4091-4102. doi: https://doi.org/10.1074/jbc.RA118.005970 Epub 2019 Jan 11.
4. Graber TG, Maroto R, Fry CS, Brightwell CR, Rasmussen BB. Measuring exercise capacity and physical function assessment in adult and older mice. J Gerontol A Biol Sci Med Sci. 2020. doi:10.1093/gerona/glaa205
5. Graber TG, Rawls BL, Tian B, Durham WJ, Brightwell CR, Brasier AR, Rasmussen BB, Fry CS. Repetitive TLR-3-mediated Lung Damage Induces Skeletal Muscle Adaptations and Cachexia. Exp Gerontol. 2018;pii: S0531-5565(17)30667-8. doi: 10.1016/j.exger.2018.02.002.
